# Unveiling the Anti-Adhesive Potential of Tetraspanin CD9 Peptides against *Pseudomonas aeruginosa* in Human Keratinocytes

**DOI:** 10.1101/2023.10.23.563549

**Authors:** Khairiyah Murad, Sharaniza Ab-Rahim, Hassanain Al-Talib

## Abstract

Multidrug-resistant *P. aeruginosa* strains are becoming a public health problem worldwide, causing numerous nosocomial infections. Adhesion of bacteria to host cells is a crucial step in infection, hence interruption of this stage can reduce bacterial infection. Tetraspanin CD9 was chosen for this study as it has been implicated in the pathogenesis of bacterial infections in a previous study. The aim of this study is to investigate the adhesion inhibition of tetraspanin CD9 peptides against *P. aeruginosa* in human keratinocytes. HaCaT cells were infected with *P. aeruginosa*, prior to treatment with CD9 peptides. The CD9 peptides cytotoxicity testing was determined by MTT assay. Bacterial adhesion was also determined quantitatively by counting viable bacterial cells and qualitatively by Giemsa staining and transmission electron microscope. Inflammatory markers (IL-8 and IL-6) expression was measured by Elisa assay. CD9 peptides did not affect HaCaT cell viability and inflammatory markers release. This study successfully demonstrated that CD9 peptides reduced *P. aeruginosa* adherence. Colonies produced by *P. aeruginosa* isolates treated with CD9 peptides were significantly reduced. Giemsa staining and TEM showed that treated samples had lower bacterial density and were located farther from the cells. These data suggest that tetraspanin CD9 peptides as the potential therapeutic approach against *P. aeruginosa* due to its property that inhibits bacterial adhesion without killing the bacteria, whereby at the same time does not adversely affect the nature of host cells.

## Introduction

*Pseudomonas aeruginosa* (*P. aeruginosa*) is the most common gram-negative bacterium causing nosocomial infections, especially in cystic fibrosis patients, burn and wound patients (1). *P. aeruginosa* is normally isolated in soil, contaminated and moist surfaces such as the bottles, sinks, medical equipment like catheters, and food trays (2). *P. aeruginosa* typically exists in the form of biofilms, attached to the surfaces of the host cells or in form of planktonic, as a unicellular microbe that moves freely with its flagellum (3). Reports have shown that *P. aeruginosa* is resistant to β-lactams, carbapenems, aminoglycosides, and fluoroquinolones antibiotics, which limits the effective choice of treatment options against *P. aeruginosa* infections (4).

According to Bassetti et.al (2018), the mortality rate among patients with carbapenems-resistant Pseudomonas infections has increased to 71% due to limited effective treatment (5). Recent report presented that multidrug resistant *P. aeruginosa* (MDR-*P. aeruginosa*) has been the common cause of superinfections such as ventilator-associated pneumonia and bloodstream infections in critical COVID-19 patients (6). There is an increased trend of COVID-19 and pneumonia, and an increase in *P. aeruginosa* bacteraemia rates has been observed during the COVID-19 pandemic period (7).

Adhesion of bacteria to the host cells is the key step to infection progression. Adhesion of *P. aeruginosa* to the surface of human cells is mediated by adhesins such as flagella, type IV pili, and quorum sensing mechanisms that facilitate the development of biofilm formation, and allow *P. aeruginosa* to attach and move around the surfaces of the host cells (8). Once bacteria colonise host cells, it is difficult to eradicate them, especially if biofilm develops. Bacterial attachment to the surface of the host cell can be inhibited by disrupting the binding surface of bacteria on the host cells. Inhibiting bacterial adhesion to host skin cells is beneficial because this mechanism acts as an indirect bacterial kill that does not affect the nature of our skin cells. With increasing concerns about the rising rates of MDR and *P. aeruginosa* nosocomial infections, several new alternative agents are urgently needed to reduce the extent of resistance and infections. The development of new antibiotics is limited, costly, and time-consuming, and sooner or later, *P. aeruginosa* will develop resistance to them (9). Therefore, new treatment options are urgently needed worldwide.

Recently, the development of anti-adherence and antimicrobial peptides has been critical for targeting multi-drug resistance (MDR) isolates, with a focus on low toxicity and low risk of bacterial resistance (10–12). Over the past decade, some pioneering studies on recombinant or synthetic peptides and/or tetraspanin antibodies as effective bacterial adhesion inhibitors have shown promise in combating various gram-negative and positive bacteria as well as viruses, although they are still far from application in healthcare settings such as against *Staphylococcus aureus, Salmonella typhimirium, Neisseria meningitidis* and *Escherichia coli* on the respective human cells (13–16). Our team reported antimicrobial effects of CD9 peptides against four bacteria that cause underarm odour, including *Micrococcus luteus, Bacillus subtilis, Staphylococcus epidermidis*, and *Corynebacterium xerosis* (17). This has evidently shown that tetraspanin-based treatment can disrupt several bacterial receptor proteins, responsible for the binding of pathogens to human cells (18). Due to the unique mode of action, there is a low risk of development of bacterial resistance than with conventional antibiotics.

Tetraspanins are a superfamily of transmembrane proteins with 33 members, consisting of four transmembrane domains (19). The unique structure of tetraspanins, which contains specific α-helices and small residues such as Gly and Ala in the transmembrane region, exhibits high variability that allows protein dynamics, helix-helix interactions, homo- and heterodimerization to form the tetraspanin network or tetraspanin microdomains (20). Given the broad tissue distribution and expression, tetraspanins have been discovered to be involved as molecular organisers within the extracellular loop region for viral entry and bacterial adhesion to host cells by interacting with bacterial cells via receptors for bacterial adhesins, subsequently leading to the pathogenesis (13,21,22). In the past, a tetraspanin-based treatment has been patented for the treatment of cancer, allergic diseases, and anaphylaxis (23,24).

However, to date, there is one limited study investigating the role of tetraspanin CD9 on *P. aeruginosa* (25,26). The tetraspanin CD9 was selected for this study because it has been implicated in the pathogenesis of bacterial infections in a previous study. In Malaysia, this is the pioneering investigation on the role of tetraspanins in bacterial infections. Earlier to the current study, our team has reported the direct antimicrobial effects of CD9 peptides which include anti-biofilm activities against *P. aeruginosa* (27). This project is in line with one of the strategies of the Ministry of Health (development of the Malaysian Nosocomial Infection Surveillance System), to streamline the current surveillance method for better management of patients with MDR infections and improve the quality of life.

Therefore, this study aims to further demonstrate the role of tetraspanin CD9 peptides as an alternative strategy to attenuate bacterial adherence to human keratinocytes without the risk of developing resistance, since CD9 peptides are considered host proteins. In the current study, we provided qualitative results on the anti-adherence properties of CD9 peptides against *P. aeruginosa* through Giemsa staining and transmission electron microscopy (TEM) which were not presented in the previous study. We hypothesised that the CD9 peptides will inhibit the adherence of *P. aeruginosa* to human keratinocytes without affecting host cell metabolic activity. This study provided the potential for future large-scale work that can be formulated and marketed to be used as an alternative to the currently used antibiotics and as a more targeted treatment strategy for infections caused by *P. aeruginosa*, perhaps leading to a better quality of treatment for *P. aeruginosa* without resistance development and reduce the economic burden on the healthcare system in combating MDR isolates.

## Methodology

### Bacterial strain

Two *P. aeruginosa* isolates were used in this study to compare the effects of CD9 peptides, which are a *P. aeruginosa* reference strain (Schroeter) Migula (ATCC 27853) and a clinical sample-isolated MDR-*P. aeruginosa* isolate (resistant to Ciprofloxacin, Ceftazidime and Gentamicin). Both isolates were provided by CPDRL, Hospital Al-Sultan Abdullah, Malaysia. The profile of bacterial resistance was also provided from the databases of CPDRL. For analysis, bacterial colonies were inoculated in Luria Bertani broth (LB, Oxoid, UK) broth and the turbidity was adjusted to 0.5 McFarland’s standard or 1.0x10^8^ CFU/mL by using a McFarland densitometer (Biomerieux, USA).

### Tetraspanin CD9 Peptides

Tetraspanin CD9 was selected for this study due to the high expression levels in keratinocytes and the involvement in the pathogenesis of bacterial infections in previous studies (15,21). Tetraspanin CD9 peptides were derived from the primary sequence of the large extracellular (EC2) domain of CD9, represented by a 15 amino acids sequence (EPQRETLKAIHYALN) with tetramethyl rhodamine (TMR)-tagging, as used in the pioneering work of Ventress et al., (2016). CD9 peptides were purchased from GenScript (USA) with 98% purity in lyophilized form.

### Cell Culture

The HaCaT cell line was purchased from iCell (Shanghai, China) and routinely maintained in a 25cm^2^ tissue culture flask. Cells were grown in complete growth media containing DMEM (Nacalai Tesque, Japan) supplemented with 10% foetal bovine serum (FBS, Tico Europe, Netherlands) and 1% penicillin-streptomycin (Nacalai Tesque, Japan). HaCaT cells were grown to confluent at 37ºC in humidified 5% CO_2_. Cells were harvested by trypsinization (Nacalai Tesque, Japan) and centrifuged at 1000 g for 5 mins upon reaching 80% confluency. The cell pellet was resuspended in cell culture media without antibiotics for further analysis.

### Cytotoxicity Testing of CD9 Peptides on HaCaT Cells

This experiment was performed to ensure that CD9 peptides treatment had no adverse effects on HaCaT cells. The HaCaT cells were grown in a 96-well plate at 1.5x10^5^ cells per well (100 µL/well) for 24 hours and treated with CD9 peptides (0.5 µM) and 0.1% DMSO alone at 37ºC in the presence of 5% CO_2_ for another 24 hours. The untreated cells were supplemented with DMEM and 10% FBS only. Cell viability assay was carried out following the protocol of the kit by Nacalai Tesque, Japan. Briefly, after 24 hours of incubation, 10 µL of MTT solution was added and incubated at 37ºC with 5% CO_2_ for 4 hours. Then, the contents of the wells were then removed and 100 µL of MTT solubilisation solution was added to dissolve the crystal formazan formation. Absorbance values were measured at 570nm (Tecan, Switzerland).

### Bacterial Adherence Assay

HaCaT cells were grown in a 24-well plate at 2.0x10^5^ cells per well and incubated at 37ºC in the presence of 5% CO_2_ for up to 24 hours until 90% confluency. Cells were washed with phosphate buffer saline (PBS, 1st BASE, Singapore). In treated wells, cells were infected with 500 µL of bacterial suspension in DMEM supplemented with 10% FBS (adjusted to 0.5 Mc Farland), followed by the addition of 500 µL of CD9 peptides (0.5 µM). Meanwhile, the untreated samples were added with DMEM supplemented with 10% FBS only. Cells were incubated at 37ºC for 3 hours in the presence of 5% CO_2_. The cells were washed three times with PBS to remove the non-adherent bacteria.

The cells were detached with trypsin for 10 minutes and then 900 µL of LB broth was added. Ten-fold serial dilutions (1:10 to 1:100,000) of *P. aeruginosa* suspension in fresh LB broth were performed and 100 uL from three dilutions (1:1000, 1:10,000 and 1:100,000) were plated on Mueller Hinton agar (Oxoid, UK). The plates were incubated overnight at 37ºC, and colonies on the plates were counted. Bacterial adhesion was expressed by colony forming unit per millilitre (CFU/mL) on agar plates. Only plates with 10-300 colonies were counted.

### Morphology of Bacterial Adherence

Giemsa staining and transmission electron microscope (TEM) was performed to visualise the adherence and morphology of *P. aeruginosa* to the HaCaT cells. Bacterial colonies were subcultured by inoculating them overnight in cell culture media DMEM supplemented with 10% FBS and the turbidity was adjusted to 0.5 McFarland’s standard. HaCaT cells were seeded into a 24-well microtiter plate at 2.0x10^5^ cells/mL and incubated at 37°C in the presence of 5% CO_2_ for 24 hours until they were 90% confluent. Then, the cells were infected with 500 µL of the bacterial suspension with the addition of 500 µL of CD9 peptides (0.5 µM) for 3 hours. For the untreated samples, only DMEM supplemented with 10% FBS was added to the cells.

For Giemsa staining, infected cells were washed with 1X PBS and cells were fixed with 100% methanol (Merck, USA) for 5 minutes. The fixed cells were stained with 3% Giemsa (J.T. Baker, USA) for 30 minutes, washed three times with 1X PBS and observed at X1000 magnification oil immersion (Olympus BX53, Japan). For analysis TEM, samples were cut ultra-thin and stained with uranyl and lead acetate staining, which was performed at the Faculty of Pharmacy, Universiti Teknologi MARA, Puncak Alam Campus, Selangor, Malaysia. TEM observation (JEOL JEM-2100F Field Emission TEM, Japan), operated at 200 kV, was performed by the professional staff at the Institute of Biosciences, Universiti Putra Malaysia.

### ELISA assay of IL-8 and IL-6

The cells were infected and detached following the steps in the previous section. The supernatant was collected and kept at -80°C until further use. The ELISA kit (Elabscience, USA) was used according to the manufacturer’s protocol to measure the expression of IL-6 and IL-8 in the media harvested from the infection of HaCaT cells (Elabscience, 2022, USA)

### Statistical analysis

All experiments were performed at least in triplicate in three independent experiments. Statistical Package for the Social Sciences (SPSS, Chicago, US) version 28.0 and Fiji ImageJ (US) was used for the statistical analysis and image observation respectively. Data represent the mean of three replicates ± standard deviation (SD).

## Results

### Effect of Tetraspanin CD9 Peptides on Human Keratinocytes Viability

The MTT test was also performed using only 0.1% DMSO as a control to CD9 peptides since CD9 peptides was initially dissolved in a little amount of DMSO. This was to ensure that the DMSO content in the CD9 peptides did not affect the efficacy of the CD9 peptides in the treated cells. The cell viability of HaCaT is shown in Figure 1. No changes in the cell viability and cell morphology (Giemsa stain and TEM photographs) were observed compared with the untreated group (control).

**Figure 1.**
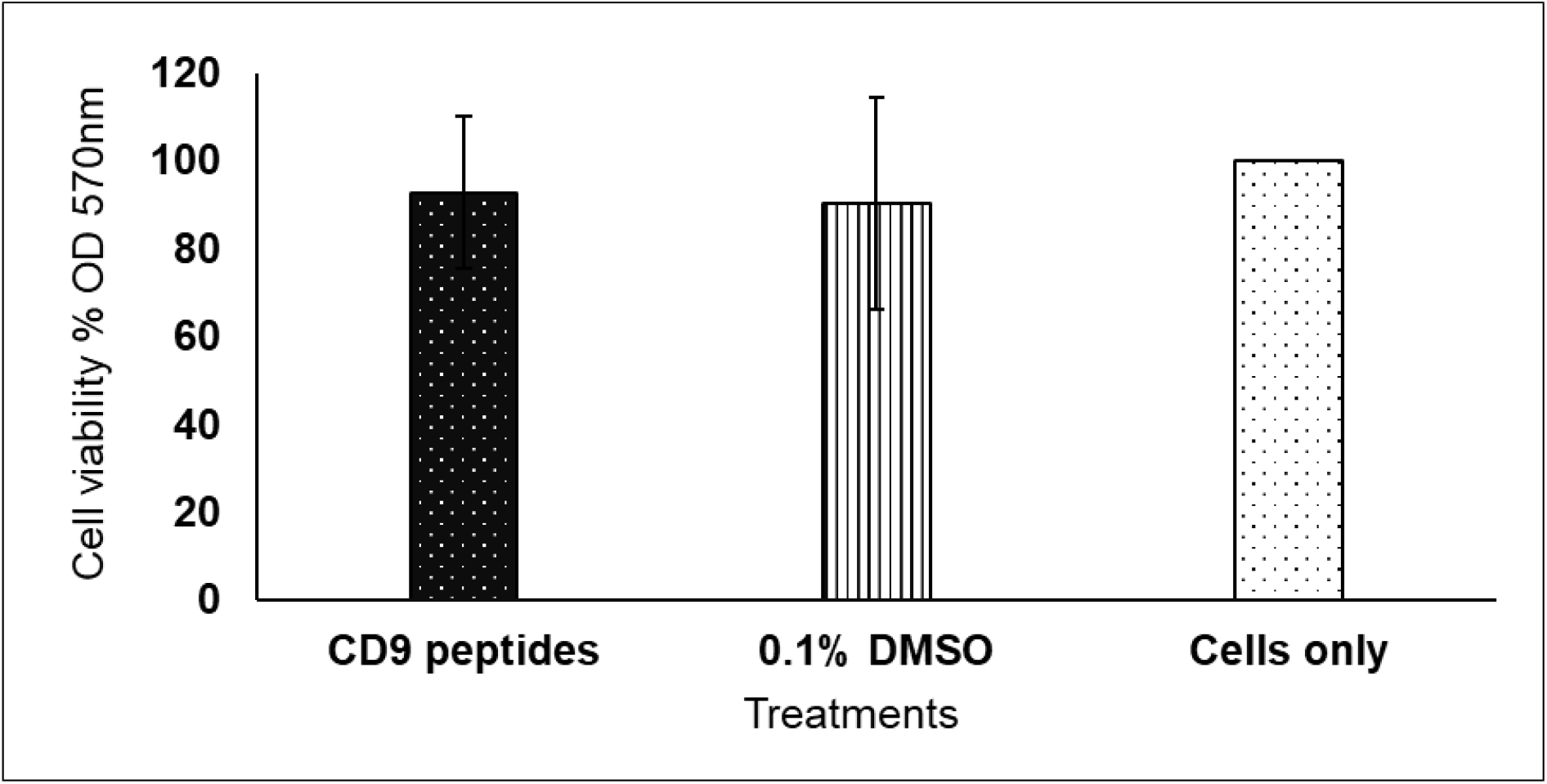
Cell viability of HaCaT cells after 24 hours of treatment with CD9 peptides and 0.1% DMSO alone. Analysis of treated cells was compared with that of control cells. No significant difference was observed in all three groups (*P* > 0.05). Absorbance values were measured at 570nm. Results are expressed as mean ± standard deviation of three readings in three independent experiments

Using the MTT dye reduction assay, HaCaT cells treated with CD9 peptides (0.5 µM) dissolved in DMSO showed no significant effect with 92.7 ± 17.4% of cells being viable after 24 hours of treatment compared to control cells, with a *P-value* of 0.48 (*P* > 0.05). The analysis showed that 0.1% DMSO alone was also tolerated by HaCaT cells, with cell viability of 90.3 ± 24.0% after 24 hours of treatment. The analysis between within three groups showed non-significant changes in the cell viability with a *P-value* of 0.282 (*P* > 0.05). Overall, this showed that CD9 peptides with the DMSO as the solvent did not cause toxicity to the cells.

### Effect of Tetraspanin CD9 Peptides on the Adherence of *P. aeruginosa* to Human Keratinocytes

The adhesion rate of *P. aeruginosa* to the HaCaT cells was determined by counting the viable cells of bacteria adhering to the cell surface. Bacterial adhesion to HaCaT cells was expressed as the mean colony-forming unit per millilitre (CFU/mL) of adherent bacteria, and the percentage of adherence inhibition was also determined. Figure 2 shows the transformed bacterial adherence rate data (log^10^ CFU/mL).

**Figure 2.**
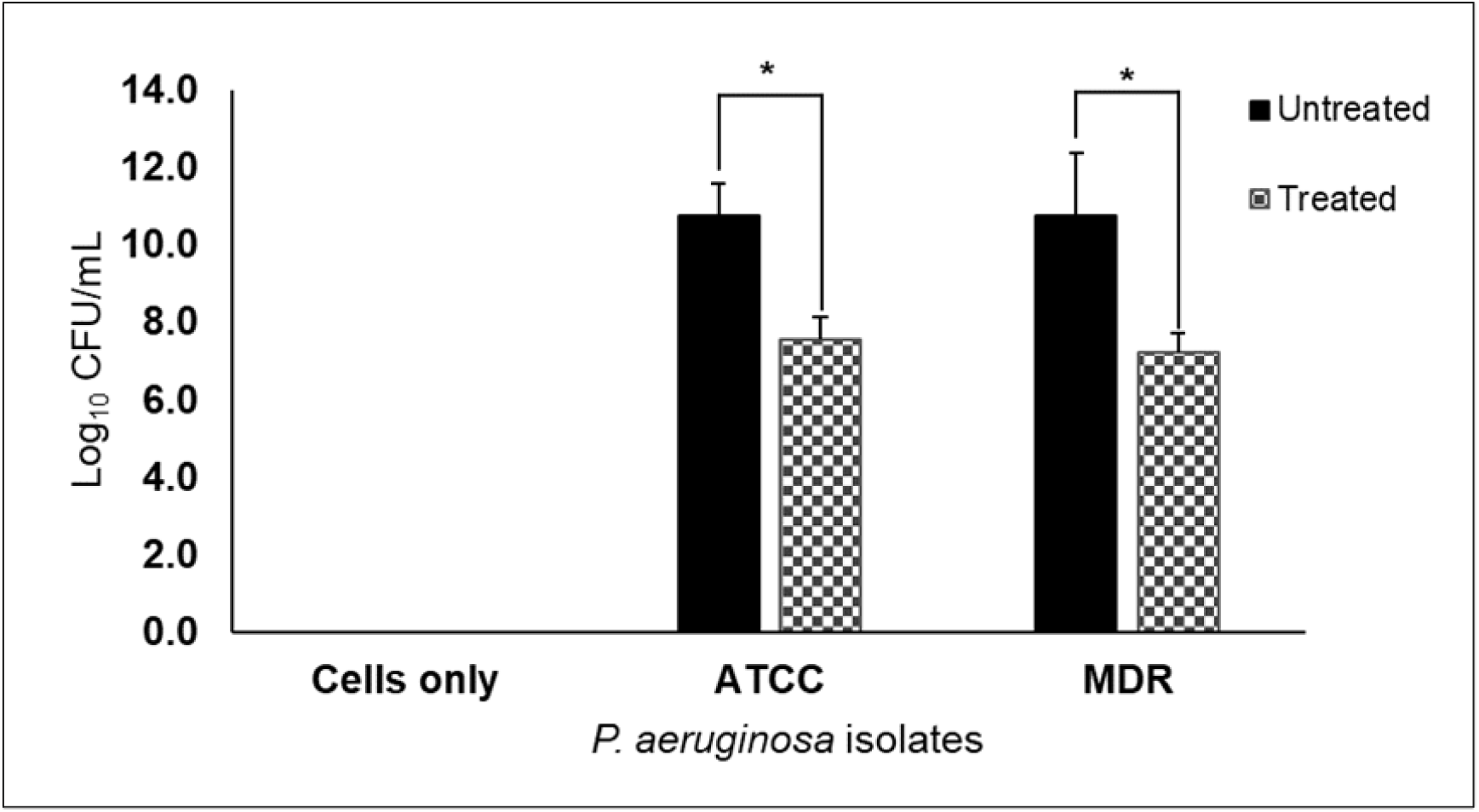
Colony-forming unit (CFU) of the adherent *P. aeruginosa* isolates treated with CD9 peptides and untreated. All analysed data were transformed into a log^10^ of CFU/mL. Results are expressed as mean ± standard deviation of three readings from three independent experiments. ^*^*P* < 0.01

In HaCaT cells infected with *P. aeruginosa* reference strain (ATCC 27853), the treated isolates had an average adherence rate of 3.41x10^7^ CFU/mL, a significant decrease compared to the untreated cells with 5.64x10^10^ CFU/mL. In HaCaT cells infected with MDR-*P. aeruginosa* treated with CD9 peptides, the rate of bacterial adherence was 1.72x10^7^ compared to the untreated cells at 5.46x10^10^ CFU/mL. These results showed a statistically significant difference at a *P* < 0.01 in colonies formed by both *P. aeruginosa* isolates treated with CD9 peptides compared with the untreated samples. In addition, the adherence rate was significantly reduced by up to 90% in both isolates treated with CD9 peptides compared with untreated cells.

### Effect of Tetraspanin CD9 Peptides on Morphology of *P. aeruginosa* and HaCaT Cells

Giemsa staining and transmission electron microscopy (TEM) were performed to visualise and identify the morphology of HaCaT cells infected with *P. aeruginosa* isolates treated and not treated with CD9 peptides. Interestingly, TEM was performed to show the internal view of the morphology of HaCaT cells and bacteria by randomly oriented cells in the dissected ultrathin sections. Observations were made to evaluate the location and density of *P. aeruginosa* in under-treated and untreated conditions. Analysis of the size of cells and bacteria, as well as the distance between cells, was performed with ImageJ based on the scale bars.

As shown in Figure 3, Giemsa staining revealed that the cytoplasm of HaCaT cells was stained bluish purple, while the nuclei were stained dark blue. *P. aeruginosa* was seen in dark purple small rods surrounding the HaCaT cells. The stained cells treated with CD9 peptides showed clear cell pores and nuclei, which was not the case for the untreated cells. Figure 4 shows the TEM photos, which show that the bacteria have a normal bacilli shape, and with some in a circular shape. This could be due to the transverse and longitudinal sections through the *P. aeruginosa* bacillus during the preparation. Interestingly, *P. aeruginosa* replication (binary fission) was observed outside the HaCaT cells by TEM.

**Figure 3.**
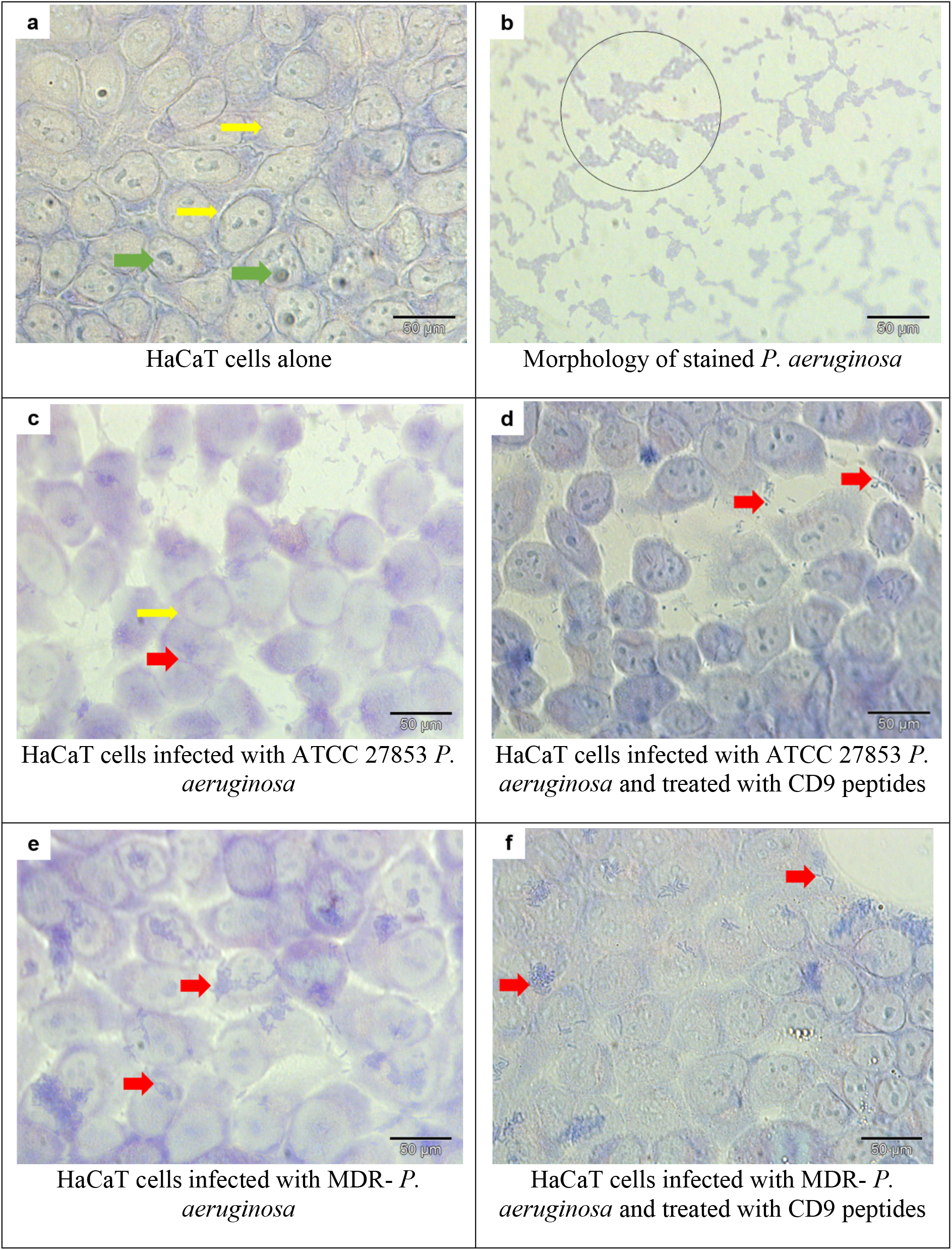
Giemsa staining at 1000X magnification and oil immersion. Red arrow: bacteria, yellow arrow: HaCaT cell membrane, green arrow: nucleus of HaCaT cells

**Figure 4.**
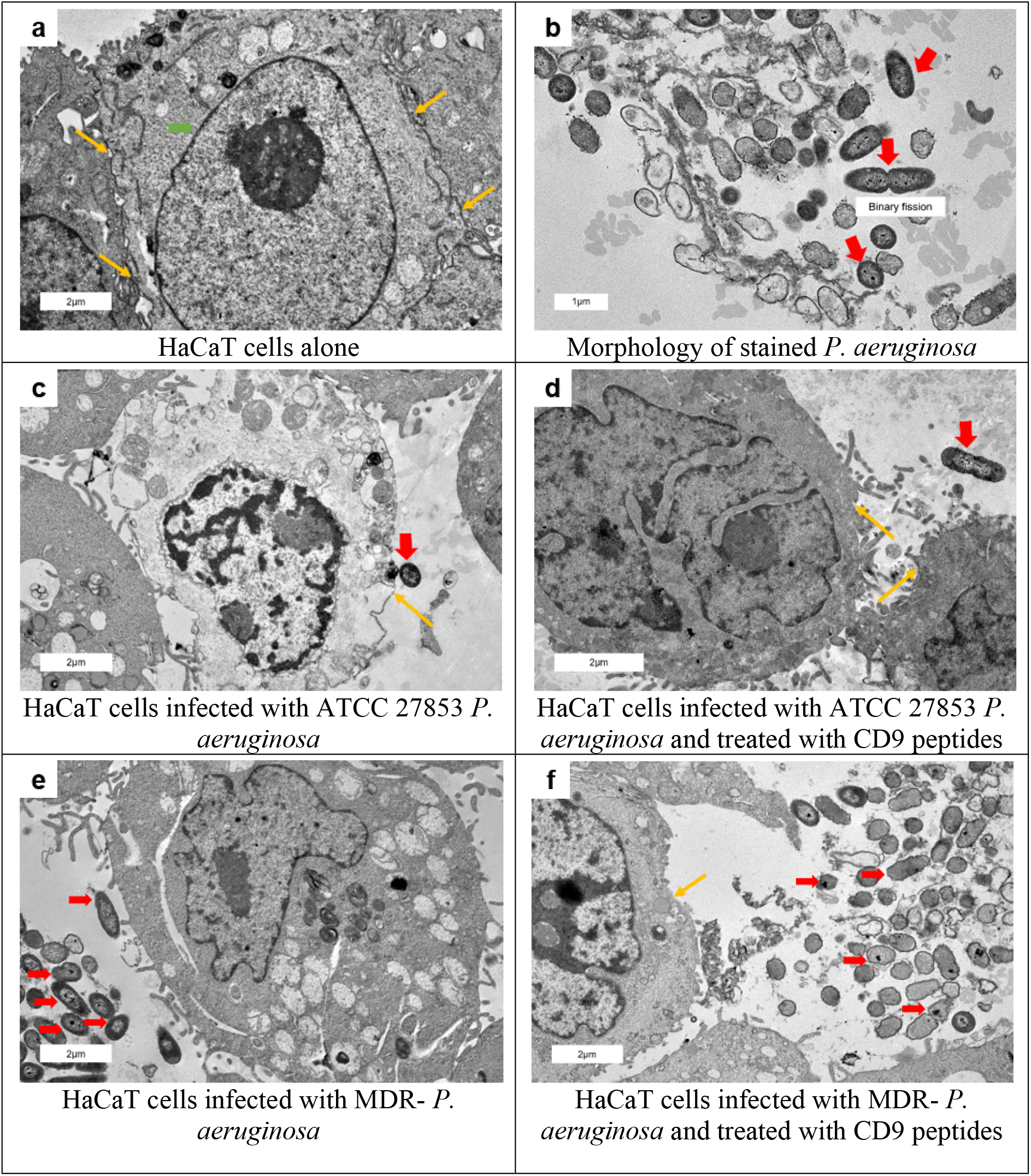
Transmission electron microscopy (TEM) photomigrograph of *P. aeruginosa* and HacaT Cells. Photomigrograph were viewed at 200 kV with the magnification of X2000 (a, c, d, e and f) and X3000 (b). Green arrow: HaCaT cells, red arrow: *P. aeruginosa*, yellow arrow: HaCaT cell membrane

Qualitative analysis showed a significant reduction in bacterial density in the cells treated with CD9 peptides compared with untreated cells. The results showed a decrease in the adherence of *P. aeruginosa* to the HaCaT cells treated with CD9 peptides compared with the untreated cells. In addition, *P. aeruginosa* were distanced away from the cells in the treated condition in comparison to the untreated. However, analysis of Giemsa staining and images from TEM using ImageJ showed there were no significant changes in the size of infected HaCaT cells compared to control cells. Analysis between treated and untreated cells also revealed no significant difference in the size of HaCaT cells. Overall, these results are consistent with the MTT assays and indicated that the CD9 peptides are not toxic to the cells.

### Effect of Tetraspanin CD9 Peptides on Inflammatory Markers Expression

Pro-inflammatory cytokines such as IL-8 and IL-6 play an important role in our immune system to clear bacteria during infection. Supernatants were obtained from HaCaT cells plated out at the same density of 2.0x10^5^ cells and infected for 3 hours with *P. aeruginosa* treated or not with CD9 peptides. The amount of interleukin released under different conditions was measured using the standard curve of IL-8 and IL-6 with R^2^ > 0.9. Figure 5 shows the measured amounts of IL-8 and IL-6.

**Figure 5.**
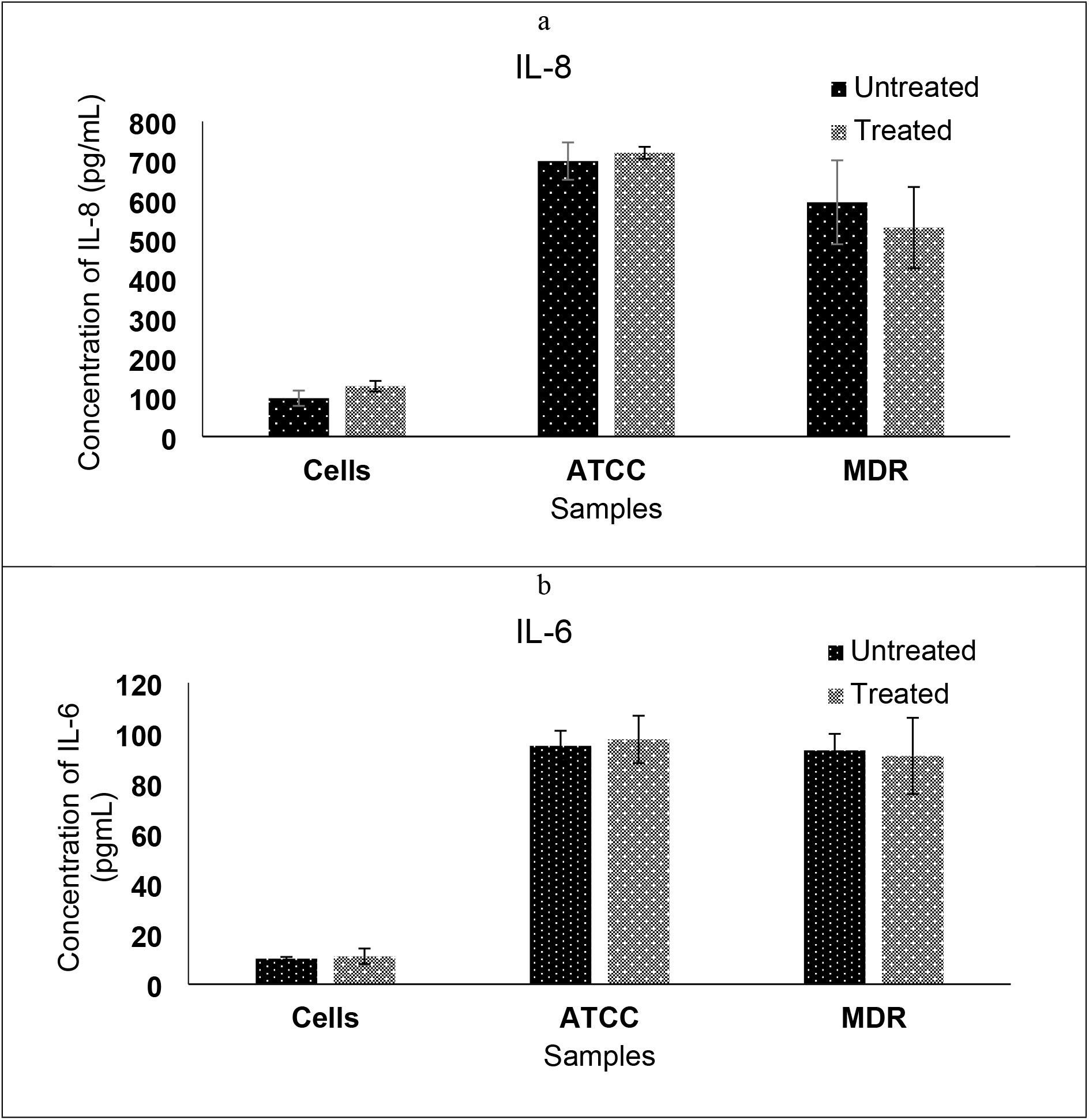
Expression of (a) IL-8 and (b) IL-6 by HaCaT cells infected with *P. aeruginosa* isolates treated with CD9 peptides and those not treated, measured by ELISA in supernatants. Results are expressed as the mean ± standard deviation of duplicates in three independent experiments

Control cells treated with CD9 peptides showed a non-significant increase in IL-8 protein secretion (127.8 ± 19.4 pg/mL) compared with untreated control cells (97.3 ± 13.4 pg/mL). These results are also consistent with the IL-6 response after the addition of CD9 peptides (11.1 ± 3.10 pg/mL) compared with cells alone (10.2 ± 0.70 pg/mL). Analysis showed that treatment of control cells with CD9 peptides did not significantly alter the levels of IL-8 and IL-6 with a *P* > 0.05. Moreover, as expected, the levels of IL-8 and IL-6 were significantly increased after infection with *P. aeruginosa* compared with control cells (*P* < 0.05).

The analysis showed that inflammatory markers were not significantly altered in the treated standard *P. aeruginosa* strain compared to the untreated conditions for the release of IL-8 and IL-6 (*P* > 0.05). IL-8 protein secretion of the treated standard *P. aeruginosa* strain was 720.2 ± 15.2 pg/mL and 97.4 ± 9.42 pg/mL for IL-6 compared to the untreated cells with 699.2 ± 47.4 pg/mL and 94.9 ± 6.01 pg/mL in IL-8 and IL-6, respectively.

However, in contrast to the standard strain, a minimal reduction in interleukin release was observed in treated MDR-*P. aeruginosa*. Treated MDR-*P. aeruginosa* had a release of 531.3 ± 103.2 pg/mL IL-8 compared to untreated cells 595.4 ± 106.5 pg/mL and 90.9 ± 15.2 pg/mL IL-6 protein secretion compared to untreated cells with 93.1 ± 6.58 pg/mL (*P* > 0.05). In general, cells treated with CD9 peptides showed no significant differences in the levels of IL-8 and IL-6 compared to untreated cells.

## Discussion

The occurrence of MDR-*P. aeruginosa* isolates has been frequently reported worldwide, and *P. aeruginosa* is known to be the major cause of wound chronic infection (28). Research on infection continues to expand the potential new antipseudomonal with the goal of reducing the incidence of MDR. Therefore, in our study, we investigated the anti-adherence activities of tetraspanin CD9 peptides against MDR-*P. aeruginosa* isolates.

The most important factor in an antimicrobial drug candidate is its safety, causing no or very low toxicity and side effects. The MTT assay was performed to ensure that the CD9 peptides did not cause harm to the host cells and did not interfere with normal cellular processes. The CD9 peptides, although dissolved in 0.1% DMSO, did not show significant negative effects on the viability of HaCaT cells. Alrahimi (2017) also reported in his dissertation that 30 minutes of treatment with 500 nM (0.5 µM) CD9 peptides did not cause a significant loss in viable cells (25). The current results are also consistent with the study by Ventress and team (2016), which showed that CD9 peptides 200 nM (0.2 µM) had no negative effects on skin cell metabolism. Furthermore, the qualitative results by Giemsa staining and TEM support these findings and suggest that CD9 peptides do not cause any cellular metabolic activity and do not alter cell morphology (Figure 3 and Figure 4). This shows that CD9 peptides do not cause toxicity when applied directly to the infected skin because the skin is composed of stratified squamous keratinized. It should be note our previous research showed that CD9 peptides has only a small direct inhibitory effect on the growth of *P. aeruginosa*, which shows CD9 peptides did not possess bacterial killing activity (27).

This phenomenon may perhaps exhibit additional benefits to humans, as this treatment boosts immunity and subsequently eliminates bacteria. In addition, a wide range of synthetic peptides is currently approved by the Food and Drug Administration (FDA) because peptides are generally characterised by high specificity, affinity, and good efficacy, and have passed the toxicity tests (29,30). Unlike antibiotics, CD9 peptides produce little to no harmful metabolites and carry a lower risk of bacterial resistance. Interestingly, *P. aeruginosa* opportunistically binds to injured epithelial cells, usually in patients with burns and wounds, more strongly than to normal cell surfaces, so anti-adhesion treatment may weaken the interaction of bacteria with the host cell receptors. The rate of bacterial adhesion to HaCaT cells was quantified by counting viable cells, while Giemsa stain and TEM produced the qualitative view of adhesion, which can differentiate the pattern of bacterial adhesion. In this study, 3% of the Giemsa stain was used to visualise the adhesion of *P. aeruginosa* to HaCaT cells. A low concentration of Giemsa stain was used to avoid false positive staining, as a study by Sungkapong and colleagues showed that a 3% Giemsa stain gave 100% sensitivity and specificity with the least unwanted dye deposition (31). In addition, analysis of TEM provides higher resolution images that can be used to analyse specific fine structures and features.

This study successfully demonstrated that CD9 peptides at a concentration of 0.5 µM significantly decreased the bacterial counts and adhered of both *P. aeruginosa* isolates to HaCaT cells compared to the untreated cells seen with Giemsa staining and TEM, supported with reduction in the colonies formed on the agar plate, demonstrating the adherence of bacteria to HaCaT cells reduced by up to 90% compared to untreated cell. Photographs obtained by Giemsa staining and TEM also confirmed the results where the treated cells had low bacterial density and some bacteria were further away from the cells compared to the untreated cells. The results clearly show the difference in bacterial distance from the cells under-treated and untreated conditions. These results are similar to those of other studies that reported the reduction of adherence of Gram-negative bacteria (N. *meningitidis, S. typhimurium* and *E. coli*) and Gram-positive bacteria (*S. aureus* and *S. pneumoniae*) to the studied cell line after treatment with tetraspanins monoclonal antibodies or/and recombinant peptides (13). Furthermore, the Giemsa staining shown in (Figure 4 d and f) that the CD9-treated cells had a clearer outline of the HaCaT membrane, and more visible pore and nucleus in comparison to the untreated cells (Figure 4 c and e). These findings may reflect the non-cytotoxic effects and protective mechanism of CD9 peptides that act as a barrier to the host cells from the destructive effects of *P. aeruginosa* infection. This situation resulted in less virulence of *P. aeruginosa* to the HaCaT cells upon treatment with CD9 peptides, resulting in less damage and maintaining the cell structures.

Nevertheless, a considerable number of *P. aeruginosa* colonies still formed on the plate. This condition was also noted by Ventress (2016), who found that despite treatment with CD9 peptides, some bacteria were found in the epidermis of the tissue-engineered skin model, but only a small amount penetrated the deeper layers. This phenomenon may indicate a reduction of bacterial load, resulting in less severe infection. This suggests that tetraspanin CD9 peptides have an indirect inhibitory effect on the attachment of bacteria to the host cells, disrupting the tetraspanin-enriched microdomain. This condition may alter the microdomain while disrupting the interaction of tetraspanins with bacterial receptors on the surface of host cells. Karam et al (2020) have previously pointed out that CD9 act as a co-receptor for the diphtheria toxin of *Corynebacterium diphtheriae*, which enhances the interaction of diphtheria toxin with its receptor and leads to infection.

These results reflect what is known about tetraspanins as indirect and nonspecific receptors for bacterial adherence, perhaps due to their properties not only as receptors but as mediators of the adhesion platform for bacterial pathogenesis(18,20,21). The CD9 peptides can exploit the properties of a receptor analogue, that competitively prevents the host CD9 receptor from interacting with the bacterial adhesins (24,32). Thus, the CD9 peptides reduced the ‘true’ bacterium-host interaction. Furthermore, a landmark study in 2011 showed that antibodies or/and recombinant tetraspanin CD63 peptides against *Neisseria meningitidis* strains lacking the gene for type iv pili only slightly reduced the bacterial adherence, compared to the wild-type strains, suggesting that tetraspanins not the direct receptor of bacterial adherence (16).

The tetraspanin CD9 is said to be involved in cell signalling and host immune response. Ventress and team (2016) reported that IL-6 and IL-8 were detected in the keratinocytes and fibroblast of the infected tissue-engineered skin. Therefore, we investigated the level of inflammatory markers (IL-6 and IL-8) in the infected cells treated with CD9 peptides. Our results showed that *P. aeruginosa* infection remarkably increased the release of interleukins. After infection, keratinocytes release many immune markers to eliminate bacteria (33). Our results are consistent with the theory reported previously, which pyocyanin production by *P. aeruginosa* can cause the release of TNF-α which induced the expression of IL-8, IL-1 and IL-6 in the infected host cells (34,35). Besides, the release of inflammatory markers helps in balancing combating the infection and minimizing host cells or tissue damage.

Interestingly, no significant changes in either interleukin level were observed after treatment with CD9 peptides compared with untreated cells. These results are consistent with the previous study reporting no effects on cytokine levels in tissue-engineered skin treated with CD9 peptides (15). The minimal changes in the release of IL-8 and IL-6 may be due to the ability of CD9 peptides to restore the physiological balance between pro- and anti-inflammatory markers during the infection. In addition, Brosseau and team (2018) have reported that CD9, CD81, and FcεRI are co-expressed and co-localized on human dendritic cells, with co-activation of CD9 and FcεRI leading to an increased release of IL-10 (36). This could be the case for the slight increase of IL-6 and IL-8 in standard *P. aeruginosa* strain, possibly CD9 peptides cross-link with other immune mediators, activating the dendritic cells, and leading to cytokine production (36). It is known that TLRs are expressed on the surface and intracellularly of the host cells to recognise the microbial components (37). The level of interleukins may have remained unchanged despite the addition of CD9 peptides to the infected cells because CD9 peptides is considered to host proteins, and therefore may not elicit any significant immune response. While there may not have been any significant changes in the interleukins release, it is possible that the CD9 peptides could provide some level of protection to the cells from further inflammation. This suggests that CD9 peptides may have a potentially beneficial effect on our immune response.

Another important point of this study showed the anti-adhesion properties of CD9 peptides may be due to the electrostatic forces introduced by the CD9 peptides, which may induce negatively charge components and slowly neutralize the surface of *P. aeruginosa*. In addition, synthetic peptides may exhibit excellent antimicrobial activity, they are relatively inexpensive and have low immunogenicity, making them favourable candidates (12,30,38). This makes the CD9 peptides very promising compared to antibiotics, which develop resistance relatively quickly.

There are some limitations in this study, including the limited synthetic tetraspanin peptide used in this study, which may mean that the results of this study are not representative of the other tetraspanin members. Therefore, it is recommended to include a different amino acid sequence of EC2-CD9 peptides as well as other peptides derived from other tetraspanin members. Moreover, this study only includes a small sample size which may not give the same antimicrobial effects of CD9 peptides to other *P. aeruginosa* strains, especially the extensively drug-resistant strains or pan drug-resistant strains. In addition, future studies should investigate the expression of the *P. aeruginosa* quorum sensing system through molecular checking, as it is the important virulence factor for bacterial biofilm formation and attachment to host cells, to better understand the anti-adherence mechanism of CD9 peptides.

## Conclusion

In conclusion, we have shown that CD9 peptides can reduce the adherence of *P. aeruginosa* to the human keratinocytes cell line without negatively affecting the cell viability and our immune response. This study successfully provides additional insights into the tetraspanin-based treatment has a dual mechanism of action as an anti-adherence agent in addition to the direct anti-biofilm effects (27). The lack of the development of resistance in microbes may be attributed to the different modes of action of synthetic peptides against bacteria compared to the standard modes of action of antibiotics. The broad specificity and low toxicity of tetraspanin-derived peptides promoted them as a good candidate for broad-spectrum antimicrobial treatment of its principle of targeting components of the host cells rather than the bacteria directly. Despite the limitations, this is a pilot study for the potential large-scale future work, formulation, and recommendation as a more targeted treatment strategy for infections caused by *P. aeruginosa*.

## Author contributions

Concepts: KM, SAR, HAT, Study design: KM, SAR, HAT, Definition of intellectual content: KM, SAR, HAT, Literature search: KM, Experimental studies: KM, Data acquisition: KM, Data analysis: KM, SAR, HAT, Statistical analysis: KM, SAR, HAT, Manuscript preparation: KM, Manuscript editing and review: KM, SAR, HAT.

## Acknowledgement

We thank Prof Dr Zolkapli Eshak from the Faculty of Pharmacy, Universiti Teknologi MARA, Puncak Alam Campus, Selangor, Malaysia for assistance in preparing the samples TEM and Mr Rafiuz Zaman Haroun from the Institute of Biosciences, Universiti Putra Malaysia in TEM viewing. Many thanks to the Institute of Medical Molecular Biotechnology (IMMB), Universiti Teknologi MARA (UiTM) for providing the facilities.

